# Effect of Live Attenuated Influenza Vaccine on Pneumococcal Carriage

**DOI:** 10.1101/343319

**Authors:** Jamie Rylance, Wouter A. A. de Steenhuijsen Piters, Sherin Pojar, Elissavet Nikolaou, Esther German, Elena Mitsi, Simon P. Jochems, Beatriz Carniel, Carla Solórzano, Jesús Reiné, Jenna F. Gritzfeld, Mei Ling J.N. Chu, Kayleigh Arp, Angela D Hyder-Wright, Helen Hill, Caz Hales, Rachel Robinson, Cath Lowe, Hugh Adler, Seher R. Zaidi, Victoria Connor, Lepa Lazarova, Katherine Piddock, India Wheeler, Emma Smith, Ben Morton, John Blakey, Hassan Burhan, Artemis Koukounari, Duolao Wang, Michael J. Mina, Stephen B. Gordon, Debby Bogaert, Neil French, Daniela M. Ferreira

**Affiliations:** Department of Clinical Sciences, Liverpool School of Tropical Medicine, Liverpool, United Kingdom; Centre for Inflammation Research, University of Edinburgh, Edinburgh, United Kingdom; Department of Paediatric Immunology and Infectious Diseases, University Medical Center Utrecht, Utrecht, The Netherlands; Department of Medical Microbiology, University Medical Center Utrecht, Utrecht, The Netherlands; Royal Liverpool and Broadgreen University Hospital, Liverpool, United Kingdom; Aintree University Hospital NHS Foundation Trust, Lower Lane, Liverpool, United Kingdo; Department of Pathology, Harvard Medical School, Boston, United States; Malawi-Liverpool Wellcome Trust Programme, Blantyre, Malawi; Institute of Infection and Global Health, University of Liverpool, Liverpool, United Kingdom

**Author notes:** both authors contributed equally to the work (Jamie Rylance, Wouter A. A. de Steenhuijsen Piters). joint senior authors (Debby Bogaert, Neil French, Daniela M. Ferreira). **Corresponding Author** Dr Jamie Rylance, Department of Clinical Sciences, Liverpool School of Tropical Medicine, Liverpool, UK, L3 5QA.

## Abstract

The widely used nasally-administered Live Attenuated Influenza Vaccine (LAIV) alters the dynamics of naturally occurring nasopharyngeal carriage of *Streptococcus pneumoniae* in animal models. Using a human experimental model (serotype 6B) we tested two hypotheses: 1) LAIV increased the density of *S. pneumoniae* in those already colonised; 2) LAIV administration promoted colonisation. Randomised, blinded administration of LAIV or nasal placebo either preceded bacterial inoculation or followed it, separated by a 3-day interval. The presence and density of *S. pneumoniae* was determined from nasal washes by bacterial culture and PCR. Overall acquisition for bacterial carriage were not altered by prior LAIV administration vs. controls (25/55 [45.5%] vs 24/62 [38.7%] respectively, p=0.46). Transient increase in acquisition was detected in LAIV recipients at day 2 (33/55 [60.0%] vs 25/62 [40.3%] in controls, p=0.03). Bacterial carriage densities were increased approximately 10-fold by day 9 in the LAIV recipients (2.82 vs 1.81 log_10_ titers, p=0.03). When immunisation followed bacterial acquisition (n=163), LAIV did not change area under the bacterial density-time curve (AUC) at day 14 by conventional microbiology (primary endpoint), but significantly reduced AUC to day 27 by PCR (p=0.03). These studies suggest that LAIV may transiently increase nasopharyngeal density of *S. pneumoniae.* Transmission effects should therefore be considered in the timing design of vaccine schedules.

**Trial registration:** The study was registered on EudraCT (2014-004634-26)

**Funding:** The study was funded by the Bill and Melinda Gates Foundation and the UK Medical Research Council.

---

Pneumonia kills more children under 5 than any other illness, and is in part preventable by immunization.^1^ In the elderly, community-acquired pneumonia is a leading cause of mortality, with *Streptococcus pneumoniae* (the pneumococcus) the most common etiologic agent^2^. The nasopharynx is colonized by the pneumococcus, and serves as the primary reservoir for human-human transmission’; this “carriage” necessarily precedes invasive infection. Influenza infection increases pneumococcal carriage density and dysregulates host immune responses. During influenza epidemics, this causes secondary bacterial pneumonia and increases mortality.^3-5^

Live Attenuated Influenza Vaccine (LAIV), a nasal spray, has been used in the United States since 2003 for influenza prevention in children and adults, and its eligibility for use for the 2018- 19 season has been confirmed by the Center for Disease Control (CDC). In the United Kingdom, annual single-dose LAIV given to children aged 2-16 years since 2013 has significantly reduced severe influenza disease.^6^ Pre-licensure and early post-marketing surveillance data showed that LAIV-protection was comparable to inactivated vaccines.^7^

Animal models of influenza infection demonstrate increased density and duration of pneumococcal carriage. Experimental human influenza infection increases carriage rates. ^3,4,8^ In mice, LAIV vaccination increases the density and duration of pneumococcal carriage, and rates of otitis media.^9,10^ In children, LAIV is associated with increased rates and density of bacterial carriage.^11^ Whilst LAIV is safe and not associated with increases in pneumococcal disease, these data suggest it could increase pneumococcal transmission to susceptible individuals.^12,13^

We used an established experimental pneumococcal challenge model to evaluate the effect of LAIV on pneumococcal carriage acquisition, density and duration. We report two trials representing distinct scenarios in the community setting: 1) *Antecedent immunisation* - individuals acquire pneumococcus following LAIV vaccination, in the period when nasopharyngeal viral replication takes place; 2) *Concurrent immunisation* - individuals already colonised with the bacteria are administered LAIV.

## Results

### Antecedent study (vaccine administered 3 days before *S. pneumoniae* inoculation)

202 participants consented, of which 142 were screened, 137 were vaccinated, 130 were inoculated, and 117 participants entered the modified ITT analysis (n=55 LAIV, n=62 control, Fig. 1B). Demographics were similar in both groups (Table S1). LAIV participants received a median 74,500 CFU/nostril of *S. pneumoniae* (range 51,000-88,000) and controls 77,250 (51,000-80,000). Mean interval between vaccination and inoculation was 3.0 days (SD 0.1). Four participants (3%) carrying pneumococcus at baseline were excluded (Table S2).

**Figure 1.**
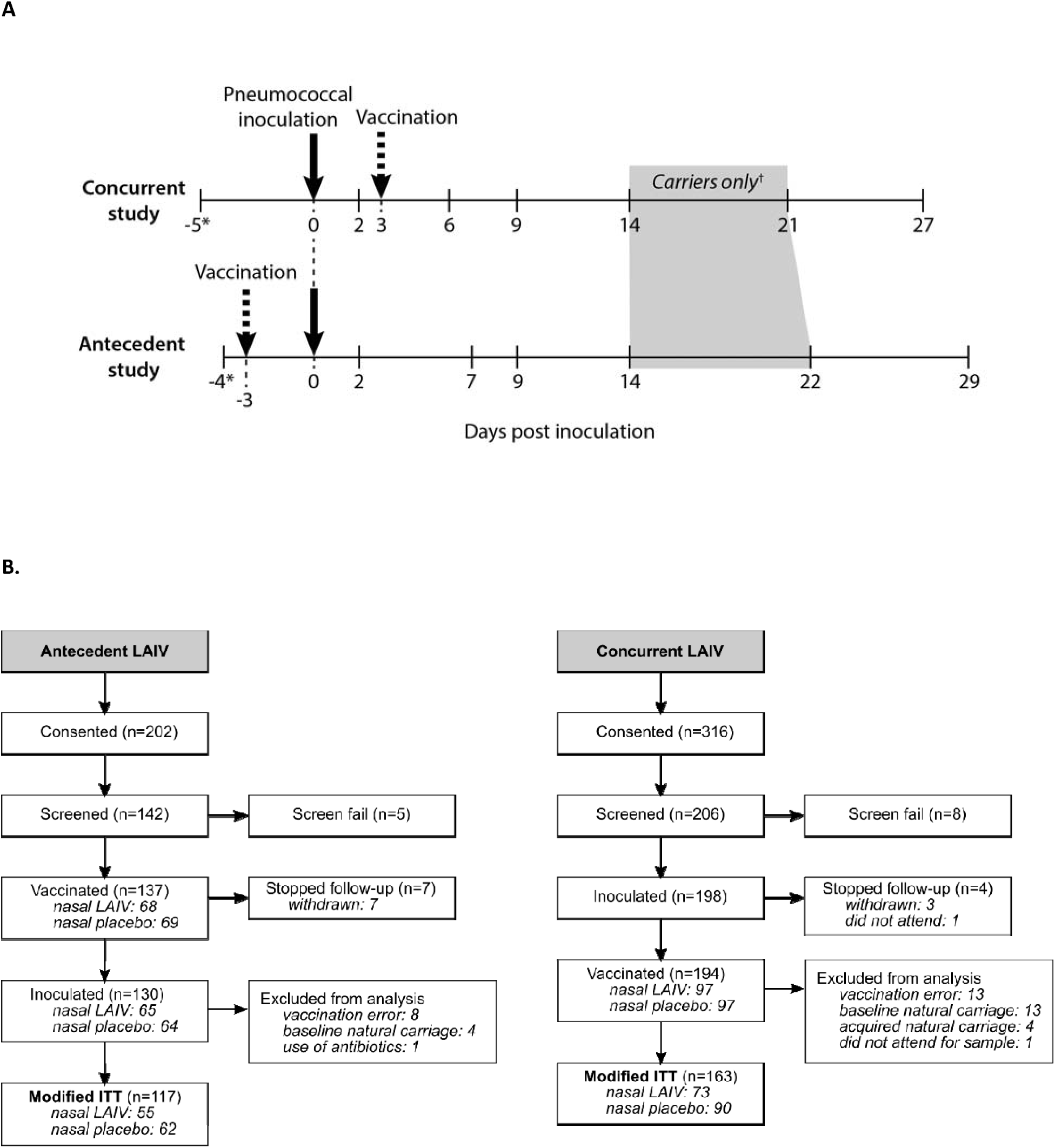
Overview study design concurrent and antecedent study and flow diagram. Panel A: Study overviews represent nasopharyngeal inoculation with *S. pneumoniae* serotype 6B as a solid arrow, and immunization as a dotted arrow. In the concurrent study, participants were screened for natural pneumococcal carriage on day -5, and for experimental pneumococcal presence on days 2, 6, 9 and 27. In the antecedent study, participants were screened for pneumococcal carriage on day -4, with further post-inoculation nasal washes on days 2, 7, 9 and 29. *represents the baseline samples relative to day of pneumococcal inoculation, with sampling performed at indicated days thereafter † For each study, only participants who had *S. pneumoniae* 6B detected by conventional microbiology on a prior sample (d14 and d21/22, grey shaded area). Samples from participants negative for pneumococcal carriage at d2, d6/d7 and/or d9 were included in the analyses and assumed negative at d14 and d21/d22. Participants entered only one study. Panel B: Participant inclusion and retention in the antecedent study (left) and concurrent study (right).

Carriage rates by conventional microbiology were consistently, though non-significantly higher in those receiving LAIV compared with control (Table 1). Comparison of the primary endpoint showed similar overall carriage rates in LAIV participants and controls (25/55 [45.5%] vs 24/62 [38.7%], OR=1.32, p=0.46). Culture-confirmed carriage at day 22 were 19/52 (36.5%) and 12/59 (20.3%) respectively (OR=2.26, p=0.06; Table 1). Molecular detection carriage rates were significantly higher in LAIV versus control at day 2 (33/55 [60.0%] vs 25/62 [40.3%], OR=2.22, p=0.03).

**Table 1.** Effect of antecedent vaccination on pneumococcal carriage status stratified by time point. Pneumococcal carriage status assessed by conventional microbiology or molecular detection *(lytA PCR)*. The number of carriage positive samples at each time point and the number of volunteers with a positive sample at any time point (i.e. carriers) according to vaccine group were reported as well as the total number included in the analysis at each/any time point. P values, odds ratios (ORs) and corresponding 95% confidence interval (CIs) were derived for each time point using a generalized estimating equations (GEE) model with time point, vaccine group and the interaction between time point and vaccine group as covariates, and participant as cluster effect.

|  | LAIV<sup>†</sup> | Control | OR (95% CI) | P value |
| --- | --- | --- | --- | --- |
| Classical microbiology |  |  |  |  |
| d2 | 23/55 (41.8%) | 20/62 (32.3%) | 1.51 (0.71-3.21) | 0.29 |
| d7 | 24/55 (43.6%) | 22/62 (35.5%) | 1.41 (0.67-2.96) | 0.37 |
| d9<sup>‡</sup> | 23/52 (44.2%) | 19/60 (31.7%) | 1.71 (0.79-3.70) | 0.17 |
| d14<sup>§</sup> | 22/54 (40.7%) | 16/61 (26.2%) | 1.93 (0.88-4.25) | 0.10 |
| d22<sup>§</sup> | 19/52 (36.5%) | 12/59 (20.3%) | 2.26 (0.97-5.27) | 0.06 |
| d29<sup>‡</sup> | 10/33 (30.3%) | 7/40 (17.5%) | 2.05 (0.68-6.18) | 0.20 |
| any time point | 25/55 (45.5%) | 24/62 (38.7%) | 1.32 (0.63-2.77) | 0.46 |
| Molecular detection (*lytA* PCR) |  |  |  |  |
| d2 | 33/55 (60.0%) | 25/62 (40.3%) | 2.22 (1.06-4.66) | 0.03 |
| d7 | 28/55 (50.9%) | 28/62 (45.2%) | 1.26 (0.61-2.61) | 0.53 |
| d9<sup>‡</sup> | 23/52 (44.2%) | 20/60 (33.3%) | 1.59 (0.74-3.41) | 0.24 |
| d14<sup>§</sup> | 23/54 (42.6%) | 18/61 (29.5%) | 1.77 (0.82-3.83) | 0.15 |
| d22<sup>§</sup> | 18/52 (34.6%) | 15/59 (25.4%) | 1.55 (0.69-3.52) | 0.29 |
| d29<sup>‡</sup> | 12/33 (36.4%) | 9/40 (22.5%) | 1.97 (0.71-5.49) | 0.20 |
| any time point | 38/55 (69.1%) | 41/62 (66.1%) | 1.14 (0.53-2.51) | 0.73 |

<sup>†</sup> Data exclude 8 participants who did not receive the required dose of the vaccine due to a systematic medication error (see text).
<sup>‡</sup> 2 Participants received antibiotics after d7: subsequent samples are treated as missing within the analysis
<sup>§</sup> d14 and d22 nasal washes were performed only in participants who had *S.pneumoniae* 6B detected by conventional microbiology on a prior sample. For the purposes of analysis, non-carriers were presumed to continue to be carriage-negative at d14 and d22.

Median duration of carriage was not different between groups by conventional microbiology (22 days [IQR 22-29] vs 22 days [IQR 14-29], p=0.09) or by molecular detection (22 days [IQR 7- 29] vs 14 days [IQR 7-22], p=0.45) for LAIV compared with control group, respectively.

Mean densities of carriage by conventional culture were consistently increased in the LAIV group, and reached statistical significance at d9 with a 10-fold (1 log_10_) increase in carriage density in the LAIV group (2.82±1.78 vs 1.81±1.39 log_10_ titers, p=0.03, Fig. 2A and Table S3). Results by molecular detection were similar, with statistically significant differences at day 2 (2.21±1.33 vs. 1.54±1.45 for LAIV and control respectively, p=0.03, Fig. 2B and Table S3). Post-hoc analysis excluding 4 individuals with confirmed concurrent viral infections (all controls, Fig S2 volunteers #20, #21, #23 and #64), demonstrated more pronounced differences, including d2 (2.18±1.33 vs 1.42±1.42, p=0.02), and d9 (2.26±1.88 vs 1.39±1.55, p=0.03; Table Se).

**Figure 2.**
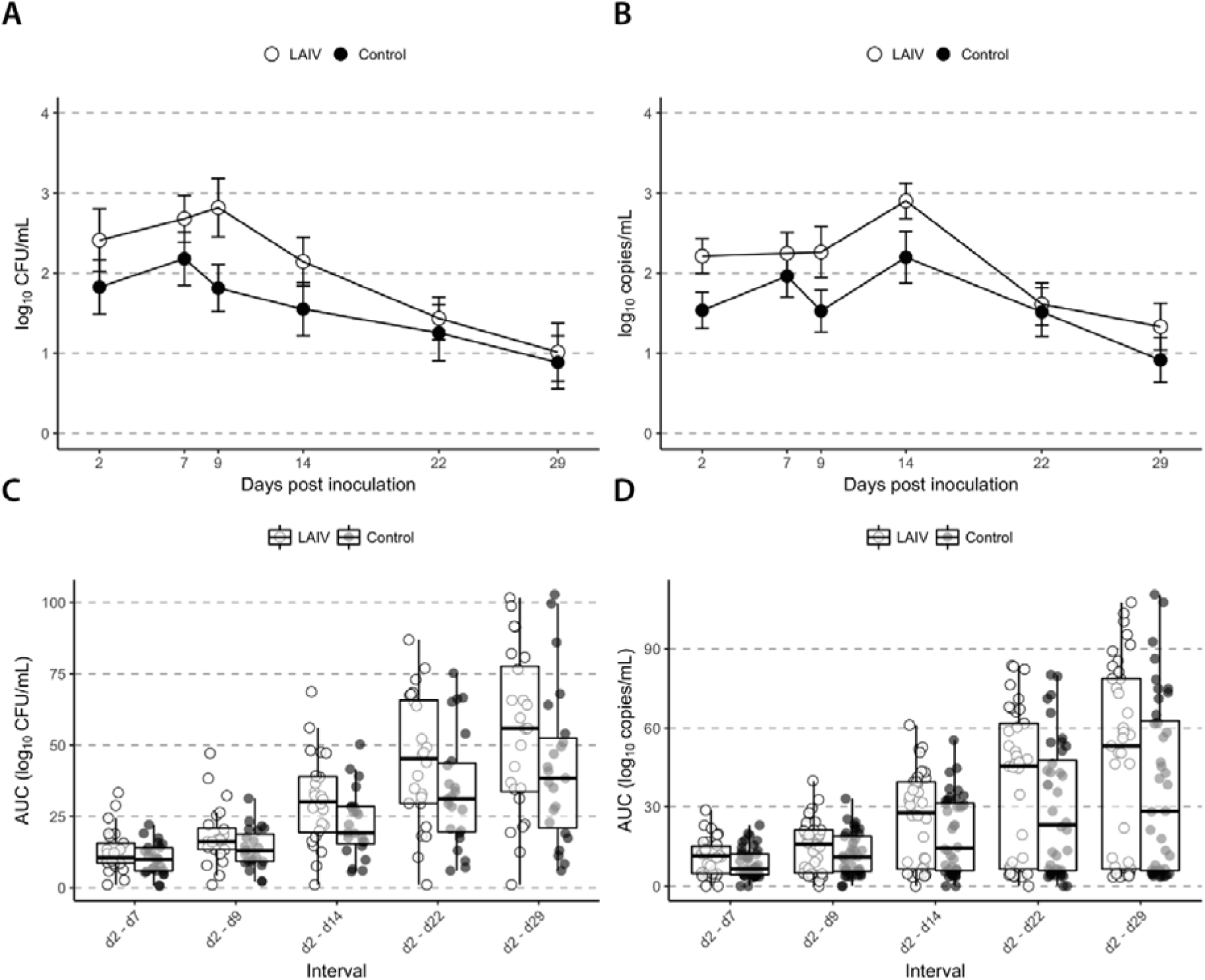
Effect of antecedent vaccination on pneumococcal colonization density dynamics. Density dynamics following pneumococcal inoculation are calculated from conventional microbiology (log_10_[CFU/ml+1], panels A and C) or molecular methods (log_10_[copies/ml+1], panels B and D). Participants receiving LAIV are represented by open circles, and controls by closed circles. Panels A and B: Mean density of *S. pneumoniae* for each nasal wash time point amongst participants in whom serotype 6B was detectable at any point. Bars represent standard errors. Panels C and D: Area under the curve of *S. pneumoniae* density from nasal washes of participants in whom serotype 6B was detectable at any time point. Boxes denote the median value (horizontal line) with 25^th^ and 75^th^ percentiles, and whiskers denote the boundary of 1.5 times the interquartile range. Data exclude 8 participants who did not receive the required dose of the vaccine due to a systematic medication error (see text).

In colonized individuals, AUC of carriage density was increased in the LAIV group versus controls by both conventional microbiology (Fig. 2C, Table S4) and molecular detection (Fig. 2D and Table S4), although statistical significance was only reached for the d2-d14 interval by conventional methods. However, amongst participants without viral coinfection, the AUC bacterial density was significantly increased between day 2 and days 7, 9 and 14 by molecular detection (p=0.03, p=0.04 and p=0.04 respectively), and between day 2 and day 14 by conventional microbiology (p=0.03, Table Sf). Individuals’ density curves are shown in Figure S2.

### Concurrent study (vaccine administered 3 days after *S. pneumoniae* colonisation)

316 participants consented, 206 were screened, 198 were inoculated and 194 vaccinated. Data from 163 participants entered the modified ITT (n=73 LAIV, n=90 control; Fig. 1B). Included participants demonstrated similar demographics in both groups (Table S5). Median inoculation doses were 82,167 CFU/nostril (range 60,667-93,000) and 81,083 (60,667-93,000) for LAIV group and control group, respectively. Data from seventeen (10%) participants were excluded due to *S. pneumoniae* carriage at baseline (n=13) or other time points (n=4) (Table S6). Colonization density prior to vaccination (at day 2) was not different (mean±SD 1.89±1.97 and 1.82±1.36 respectively, p=0.84, Table S7).

There was no significant difference in the primary endpoint of AUC carriage density by conventional microbiology to day 14, (mean±SD 20.93±14.07 and 26.10±14.41 in LAIV and controls respectively, p=0.11, Fig. 3A and C, Table 2). AUC by molecular detection were 21.61±14.02 and 25.89±14.94 for LAIV and control groups respectively (p=0.15, Fig. 3B and D, Table 2). For other intervals, the AUC of carriage density was consistently lower for LAIV than control (i.e. days 2 through 6, 9, 21, 27), although not significantly by conventional microbiology. By molecular detection there was significantly lower AUC for LAIV vs controls for the day 2-27 interval (35.76±25.83 vs 48.66±29.48 respectively, p=0.03). Individuals’ density curves are shown in Figure S3.

**Figure 3.**
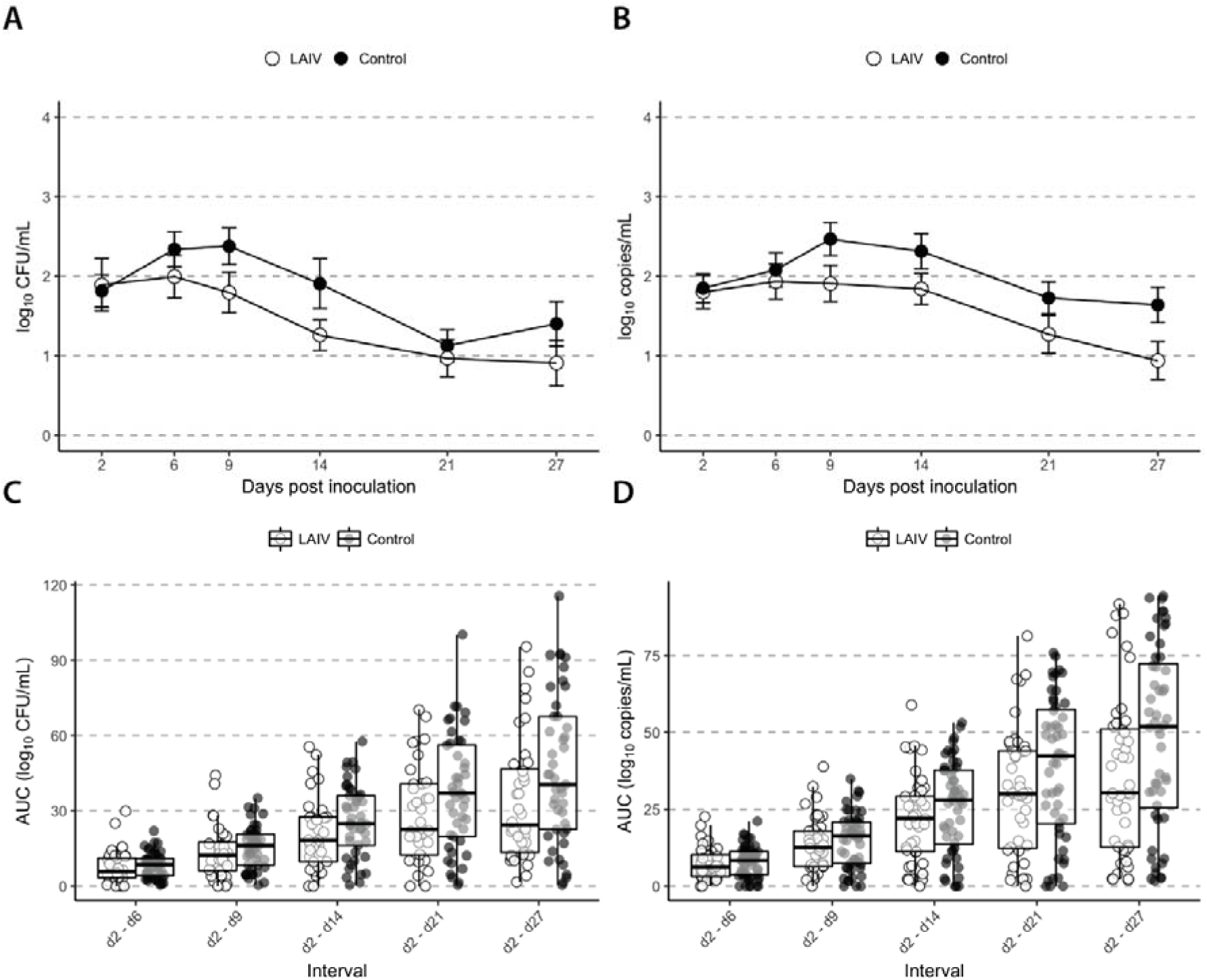
Effect of concurrent vaccination on pneumococcal colonization density dynamics. Density dynamics following pneumococcal inoculation are calculated from conventional microbiology (log_10_[CFU/ml+1], panels A and C) or molecular methods (log_10_[copies/ml+1], panels B and D). Participants receiving LAIV are represented by open circles, and controls by closed circles. Panels A and B: Mean density of *S. pneumoniae* for each nasal wash time point amongst participants in whom serotype 6B was detectable at any point. Bars represent standard errors. Panels C and D: Area under the curve of *S. pneumoniae* density from nasal washes of participants in whom serotype 6B was detectable at any time point. Boxes denote the median value (horizontal line) with 25^th^ and 75^th^ percentiles, and whiskers denote the boundary of 1.5 times the interquartile range.

**Table 2.** Effect of concurrent vaccination on the area under the density curve for each interval from day 2. Density dynamics of *S. pneumoniae* from nasal washes of participants in whom serotype 6B was detectable at any time point. Area under the curve is calculated from conventional microbiology results (log_10_[CFU/ml+1]) and molecular methods (log_10_[copies/ml+1]), using the trapezoid rule. n denotes the number of participants contributing calculable AUC values for the given period. Missing density values for time points flanked by known density values were interpolated using the mean of the known values. Sequential missing values and/or missing values at the end of a given interval were extrapolated using the average change over that interval stratified by vaccine. Differences and corresponding 95% confidence intervals (CIs) of AUCs between vaccine groups and P values were generated using generalized linear models (GLMs) with vaccine group as a covariate. AUC-differences between vaccine groups were tested in a separate GLM for each interval considered.

|  | LAIV |  | Control |  | Difference<br>(95% CI) | P value |
| --- | --- | --- | --- | --- | --- | --- |
|  | n | Mean ± SD | n | Mean ± SD |  |  |
| Conventional microbiology |  |  |  |  |  |  |
| d2 - d6 | 36 | 7.78 ± 6.58 | 45 | 8.31 ± 5.12 | 0.53 (-2.02 - 3.08) | 0.68 |
| d2 - d9 | 36 | 13.47 ± 10.21 | 45 | 15.38 ± 8.64 | 1.91 (-2.19 - 6.02) | 0.36 |
| d2 - d14 | 36 | 20.93 ± 14.07 | 45 | 26.10 ± 14.41 | 5.17 (-1.08 - 11.41) | 0.11 |
| d2 - d21 | 36 | 28.34 ± 19.05 | 45 | 37.20 ± 23.00 | 8.86 (-0.49 - 18.21) | 0.07 |
| d2 - d27 | 36 | 33.61 ± 24.46 | 45 | 45.49 ± 29.64 | 11.89 (-0.15 - 23.92) | 0.06 |
| Molecular methods |  |  |  |  |  |  |
| d2 - d6 | 44 | 7.48 ± 5.12 | 52 | 7.86 ± 5.20 | 0.39 (-1.69 - 2.46) | 0.72 |
| d2 - d9<sup>†</sup> | 44 | 13.24 ± 8.86 | 52<sup>†</sup> | 14.68 ± 9.03 | 1.44 (-2.16 - 5.03) | 0.44 |
| d2 - d14<sup>†</sup> | 44 | 21.61 ± 14.02 | 52 | 25.89 ± 14.94 | 4.28 (-1.55 - 10.11) | 0.15 |
| d2 - d21<sup>†</sup> | 44 | 30.11 ± 20.56 | 52 | 38.54 ± 22.98 | 8.42 (-0.37 - 17.22) | 0.06 |
| d2 - d27<sup>†</sup> | 44 | 35.76 ± 25.83 | 52 | 48.66 ± 29.48 | 12.91 (1.72 - 24.10) | 0.03 |

<sup>†</sup> Due to a laboratory error, molecular detection was not performed in 1 sample collected from a carrier in the control group at day 9.

Reflecting the AUC analysis, pneumococcal carriage density was consistently lower in LAIV versus controls at each time point from day 6 onwards, and was significantly different by molecular detection at day 27 (0.94±1.48 vs 1.64±1.49, p=0.03, Fig 3A and B, Table S7).

Overall rates of carriage were not different between LAIV and control groups by conventional microbiology (36/73 [49.3%] vs 45/90 [50.0%] respectively, OR=0.97, p=0.93, Table S8) or molecular detection (44/73 [60.3%] vs 52/90 [57.8%], OR=1.08, p=0.81, Table S8). We observed a decrease in carriage rates in the LAIV group compared to controls over time reaching significance at day 27 by both conventional culture (14/67 [20.9%] vs 29/82 [35.4%], OR 0.48, p=0.05) and molecular detection (14/67 [20.9%] vs 32/82 [39.0%], OR=0.41, p=0.02) (Table S8). This was consistent with lower median duration of carriage by molecular detection in the LAIV group (14 days, IQR 9-27 vs 27 days, IQR 21-27, p=0.001). Carriage duration by conventional microbiology was not different (21 days [IQR 14-27] vs 27 days [IQR 21-27], p=0.17).

There were no Serious Adverse Events related to the intervention in either study. One participant (not colonised with *S. pneumoniae*) self-administered the standby antibiotics due to upper respiratory tract symptoms.

## Discussion

We have studied for the first time the impact of a live viral vaccine on a common respiratory bacterial pathogen using a controlled human co-infection model.

Antecedent LAIV administration did not significantly alter overall pneumococcal acquisition rates, but caused significant transient effects on carriage. Increased carriage acquisition at day 2, and carriage density at days 2 through 14 were observed by molecular detection (PCR) after removal of participants with confirmed intercurrent (non-vaccine) viral co-infection.

We found no evidence of increased colonization rates, nor increased bacterial densities in those subjects already colonized when receiving LAIV. In fact, this scenario was associated with reduced carriage density and carriage rates at day 27, decreased AUC of carriage density to day 27, and earlier clearance of carriage compared with the control group.

Although our model describes colonization, the results are consistent with murine co-infection disease models of wild type influenza and pneumococcus, reinforcing that the precedence of pathogen exposure determines disease outcome: pneumococcus infection following influenza can exacerbate disease, whereas pneumococcus infection preceding influenza may reduce mortality.^14^ We hypothesise that these differences in outcome associate with different carriage densities, supported by the positive association of carriage density and pneumonia risk and severity.^13,15,16^

LAIV has been extensively safety-proven. However, murine models demonstrate that live attenuated virus replication in the nasopharynx can increase density and duration of pneumococcal carriage. A randomized controlled trial in children showed an increased pneumococcal density following LAIV.^9^ We observed similar effects: bacterial colonization density reached approximately 1 log_10_ higher post bacterial inoculation, but only when LAIV preceded pneumococcal challenge. With the order reversed, bacterial density decreased and clearance was more rapid.

Our results suggest increased pneumococcal carriage density following LAIV. Although these were not statistically significant at every time point, the antecedent study was powered to detect changes in acquisition, not density: based on previous calculations, we would require twice the number of acquisitions per arm (80 instead of 40) to detect a 2-fold difference in bacterial titres at any individual time point.^17^ For overall AUC bacterial density, post-hoc analysis suggests 50 colonized individuals per arm would be required to detect a 50% difference.

Three participants with laboratory confirmed influenza B infections (all controls) and one with rhinovirus (LAIV group), had among the highest bacterial densities of their cohorts. Given the strong association between influenza or rhinovirus infection and high bacterial carriage density, we performed a post-hoc sensitivity analysis. After removing these individuals, we detected statistically significant increases in densities in the LAIV group and, based on the AUC bacterial density analyses, a compelling pattern of increases across every time interval.

We used two complementary and consistent methods for bacterial detection; conventional microbiology and more sensitive PCR-based detection. While the latter may in some cases represent the persistence of bacterial DNA in the absence of viable pathogen, the persistence beyond one week suggests lower level colonisation (unmeasurable by conventional microbiology). Further testing of early dynamics (by PCR of serial salivary samples within the first 2 days) is underway.

We examined LAIV following pneumococcal carriage and the reverse, showing that order of events is important in colonization events and should be considered in future studies and clinical protocols. This model allows careful observation of the timing and natural history of carriage, including immunological endpoints during vaccine testing, with stable carriage rates between studies.^18-20^ Our studies reflects real-world choices made within immunisation campaigns (intramuscular vs nasal), and proxy endpoints relevant to community bacterial transmission risk during such programmes; AUC bacterial density, time-specific carriage densities, and time to clearance.^21^ Droplet transmission is a complex process, primarily determined by number of infectious particles, but sensitive to multiple factors including ventilation.^22^ Despite notable epidemics in crowded conditions,^23^ there are very limited data on inoculum dose for *S. pneumoniae*. We chose 1 log_10_ as a clinically relevant density difference; in mice, increases in colonisation density even below this threshold leads to increased transmission.^12^

Potential weaknesses of these studies were their limited size, single centre administration, and use of only one *Streptococcus pnuemoniae* serotype. We are currently assessing other serotypes within our model to address this limitation. Our model also recruited adults who, due to previous exposure to wild type influenza, are likely to have neutralizing antibodies. Any LAIV effect may be more pronounced in children with lower antibody rates and titres, increased vaccine virus reproduction and shedding, and higher rates of natural pneumococcal carriage.^24^ LAIV might lead to a transient increased density of pneumococcal carriage in adults, and this effect could be more pronounced in infants and children due to the absence of antibodies in this target group.

These studies are novel in three ways: 1) the first study in humans that directly assessed the impact of a vaccine on an unrelated human pathogen; 2) the largest conducted vaccine testing studies using a controlled human infection model; 3) the first controlled co-infection model using two live pathogens.

Future studies should investigate whether LAIV-vaccinated children are more likely to transmit nasal bacteria to their immediate contacts.^21,25^ In future vaccine protection studies, we emphasize the importance of evaluating the effect on pathogens not directly targeted by the vaccine.

## Methods

### Study design and participants

Within two single-centre, double-blinded, randomised, placebo controlled trials, we recruited two sequential cohorts: one “antecedent” cohort (LAIV was administered prior to experimental inoculation with *S. pneumoniae*), and one “concurrent” cohort (LAIV was administered following pneumococcal inoculation). Healthy non-smoking volunteers, aged 18-50, were invited to take part. Exclusion criteria were: influenza or pneumococcal vaccination or clinically confirmed disease in the preceding two years; close contact with “high-risk” individuals (children under 5, immunosuppressed, elderly); current febrile illness; use of antibiotics or immune-modulating medication; pregnancy. Historical medical data were confirmed from patient records held by the General Practitioner.

### Randomisation and masking

Using a permuted-block algorithm (1:1, blocks of 10) held in sealed envelopes, participants were randomised to receive either nasal LAIV (Fluenz Tetra or FluMist Tetra, AstraZeneca, UK, used interchangeably due to procurement shortages) paired with intramuscular placebo (0.5ml normal saline), or nasal placebo [control] (0.2ml normal saline) paired with intramuscular Quadrivalent Inactivated Influenza Vaccination (Fluarix Tetra, GlaxoSmithKline, UK) (see supplemental methods for flu strains). Vaccine and placebo were prepared by nurses independent of the study team in a separate room, and participants were blindfolded during administration. Laboratory and clinical study teams remained masked to vaccine allocation until study unblinding.

### Procedures

Experimental pneumococcal challenge protocol was followed as previously described,^26^ with serotype and dosing regimen choice as documented.^19^ Briefly, 80,000 colony-forming-units (CFU) of *S. pneumoniae* serotype 6B (strain BHN418) in 0.1ml solution were sprayed by laboratory pipette into each nostril, as per previous dose-ranging experiments. Batch-prepared mid-log-growth cultures in Vegetone broth (Sigma, Switzerland) were stored at -80°C, and independently tested by Public Health England for purity and antibiotic sensitivity. Participants were inoculated within 30 minutes of stock dilution. The dose was confirmed by dilution plating.

In the antecedent study (2015/16), LAIV vaccination preceded pneumococcal inoculation by 3 days. In the concurrent study (2016/17), the order was reversed (Fig. 1).

### End-Point Assessments

Individuals were “carriage positive” if any nasal wash culture following experimental challenge contained *S. pneumoniae* serotype 6B. Carriage was detected by conventional culture and quantified by serial dilution from nasal washes performed at days 2,7,9 and 29 (antecedent study), and 2,6,9 and 27 (concurrent study).^19,26-28^ For carriers, additional washes occurred at days 14 and 21/22. Serotype was confirmed by latex agglutination (Statens Serum Institute, Copenhagen, Denmark).^27^ Molecular detection by quantitative PCR targeted the pneumococcal *lytA* gene in nasal wash pellets, using a positive C_T_<40.^29^

### Safety

We recorded adverse events until 30 days after inoculation using structured questionnaire at study visits, additional reports from participant-initiated consultation on our 24-hour study phoneline, and self-report of administration of “standby” antibiotics (provided to study participants for use if immediately required). Participants carrying *S. pneumoniae* at their penultimate or ultimate clinic visits received a 3-day amoxicillin course after final sampling. An independent data and safety monitoring committee with access to unblinded data provided oversight of the safety and conduct of the trial.

### Statistical Analysis

The area under the curve (AUC) for bacterial density was calculated starting from day 2 according to the trapezoid rule using values of [log_10_ (bacterial density+1)] for each interval. Missing densities were interpolated from the mean of flanking density values. Sequential missing values, and missing values at the end of a given interval, were extrapolated using the average change over that interval stratified by vaccine group, with zero as a lower limit. Interpolation did not result in qualitative or important quantitative differences in results (sensitivity analysis assessed effects by randomly drawing the change from the respective interval’s Gaussian distribution around the mean-change, iterating over 1000 draws.)

#### Antecedent study

The primary endpoint was the proportion of carriage-positive participants detected by conventional microbiology. Secondary endpoints were; proportion of carriage at each time point, duration of carriage, density of carriage at each time point, AUC bacterial density to each time point, and each endpoint determined by molecular detection. With baseline carriage rates of 50% (expected based on prior data from our model), 73 participants in each arm were required for 80% power to detect a 50% relative increase in pneumococcal acquisition at any time point, after 10% drop-out.

#### Concurrent study

The primary endpoint was AUC bacterial density to day 14 by conventional microbiology amongst carriers only. Secondary endpoints included: duration of carriage, carriage proportion and density at each time point, AUC bacterial density to each time point, and each endpoint determined by molecular detection. Based on density data from the antecedent study control arm, we required 78 colonised participants (requiring 156 inclusions overall assuming 50% carriage) to detect a unit increase in log_10_(AUC carriage d2-d14) with 80% power, after 10% drop-out.

Generalised linear models (GLM) compared carriage rates at any time point, the duration of colonization, and AUC bacterial density over each interval. Generalised estimating equations (GEE) compared carriage rates and pneumococcal density at each time point, with participant as cluster effect and assuming an exchangeable working correlation structure. Post-hoc sensitivity analyses excluded cases of confirmed concurrent viral infections (Influenza B n=3; rhinovirus n=1) identified by symptom-triggered clinical review. Comparisons were two sided, with p<0.05 considered significant. Analyses were performed in R v3.3.3 (R Foundation). We report the planned modified intention to treat (ITT) analysis which excludes participants not receiving the vaccine or pneumococcal inoculation, those not completing the minimum required follow-up visits (day 2 and 7), and those with *S. pneumoniae* carriage of non-experimental serotype before inoculation (both studies) and following inoculation (concurrent study only).

During the concurrent study, we detected a systematic vaccine dispensing error by a single practitioner which resulted in 13 participants in the LAIV arm not receiving the required dose. Following advice from the trial steering group (TSG), after consultation with the Medicines and Healthcare-products Regulatory Agency, senior investigators were unblinded to prior allocations, and the trial was extended to allow the pre-specified number of LAIV allocations (n=73) to be reached. 1:1 allocation remained in place, as alteration might have led to effective unblinding of the team. The TSG recommended that data from all participants dispensed LAIV by the same practitioner *in both studies* should be excluded. This manuscript reports results from individuals unaffected by this error. Full data are given in the online supplement (Table Sa-Sd, Fig S1).

Written informed consent was obtained from all participants before enrolment. The study protocol was approved by the sponsors and the North West NHS Research Ethics Committee (14/NW/1460), and was performed in accordance with the principles of the Declaration of Helsinki and the International Council for Harmonisation Good Clinical Practice guidelines.

## Acknowledgements

The authors would like to thank all volunteers who participated in this study, the support from Clinical Research Unit staff, the RD&I research nurses (Royal Liverpool and Broadgreen University Hospital) and Catherine Molloy and Kelly Convey (Liverpool School of Tropical Medicine). The authors also thank the members of the Data Monitoring and Safety Committee: Prof Robert C. Read, University of Southampton (Chair); Prof Brian Faragher (LSTM); Dr Christopher Green (University of Oxford).

## Author contributions

JR, DW, SBG, NF, DMF designed the trial.

JR, ADHW, HH, CH, RR, CL, HA, SRZ, VC, LL, KP, IW, ES, BM, JB, HB conducted the trial according to the study protocol.

SP, EN, EG, EM, SPJ, BC, CS, JRe, JFG participated in site work including laboratory processing, data collection, and challenge preparation.

JR, WAASP, SP, EN, EG, EM, SPJ, BC, CS, MLC, KA, AK, DW, MJM, DB, NF, DMF contributed to laboratory analysis, data interpretation, statistical analysis, and literature search.

JR, WAASP, MJM, DMF drafted the report.

All authors contributed to critical review of the report.

## Data availability statement

Streptococcus pneumoniae BHN418 sequence is available GI:557376079 <u>(https://www.ncbi.nlm.nih.gov/nuccore/557376079).</u> The datasets generated during and analysed during the current study, and the statistical analysis plan, are available from the Dryad data repository (DOI ***) [is given by Dryad only when accepted for publication]

